# Large Scale-Free Network Organization is Likely Key for Biofilm Phase Transition

**DOI:** 10.1101/630103

**Authors:** Kumar Selvarajoo

## Abstract

Non-linear Kuramoto model has been used to study synchronized or *sync* behavior in numerous fields, however, its application in biology is scare. Here, I introduce the basic model and provide examples where large scale small-world or scale-free networks are crucial for spontaneous *sync* even for low coupling strength. This information was next checked for relevance in living systems where it is now well-known that biological networks are scale-free. Our recent transcriptome-wide data analysis of *Saccharomyces cerevisiae* biofilm showed that low to middle expressed genes are key for scale invariance in biology. Together, the current data indicate that biological network connectivity structure with low coupling strength, or expression levels, is sufficient for *sync* behavior. For biofilm regulation, it may, therefore, be necessary to investigate large scale low expression genes rather than small scale high expression genes.

## 1. Introduction

Living organisms are complex, dynamical and dissipative systems, often considered by physicists to be in a far from equilibrium state [1,2]. That is, for their survival, living systems exchange matter dynamically and are able to evolve spontaneously under certain environmental perturbation towards a critical point for a phase transition. This happens without fine-tuning their system parameters [3, 4]. The formation of biofilm by certain microorganisms, such as the bacteria *Escherichia coli* and the fungi *Saccharomyces cerevisiae*, is a common example [5, 6].

Also, on numerous occasions, living organisms display synchronized emergent behavior or response; the collective motion of fish on arrival of predators, the metabolic cycles in yeast or the circadian gene expression oscillations [7, 8]. Such phase transformation is known to break the symmetry of the system leaving it invariant, or in a collective mode [9]. At the critical point, the system achieves universality, that is, all differences between individuals will be reduced to follow common “universal” properties. Such fascinating self-organizing behavior of biology has triggered scientists across diverse disciplines to study the underlying *sync* mechanisms.

## 2. Kuramoto Model – Large scale network oscillators *sync* with less coupling strength

Over the last few decades, notable mathematicians, theoreticians and physicists have explored numerous analytical methods and have written extensively on interpreting the emergent self-organization or *sync* behavior in nature [2,10-12]. A pioneering work back in 1975, Yoshiki Kuramoto published an article on the synchronization of coupled oscillators that received worldwide attention [13]. The model shows that a dynamical system can collectively synchronize even despite differences in the individual node’s natural frequencies. The governing equation of the model is:

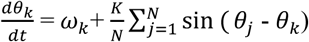

where *ω*_*k*_ is natural frequency, *θ*_*k*_ phase angles, *K* coupling strength of *N* limit-cycle oscillators. The assumptions of model include the oscillators are almost identical, their interactions depend sinusoidally on the phase difference and weak coupling between each pair of system.

Notably, Kuramoto included a clever transformation that enabled the non-linear model to be solved exactly in the limit when *N* → ∞. This required the development of the “order parameter”, that measures the synchronization between nodes:

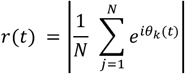

Here, *r*, or sometimes noted as *q*, represents the phase-coherence of the oscillators where *r* = 0 represents no *sync* and *r* = *e*^*iθ*^ indicate perfect *sync*.

Figure 1A shows that the Kuramoto model, in its original form, is able to synchronize in a large scale coupled network even for a specified weak coupling strength. It not only shows whether partial or complete *sync* of the network occurs (*r* = 1 is complete *sync*), but also predict the response of the network system near *sync* and the attractor region to which the system will converge. Thus, there have seen numerous variants of the Kuramoto model that have been investigated to study non-linearities in ecology, physics, social networks and biology. For instance, in electrical design, disordered Josephson junctions (two superconducting metals that are separated by a thin layer of insulator) showed partial synchronization to complete phase locking, where the transition phenomena were explained by the Kuramoto model [14].

**Figure 1.**
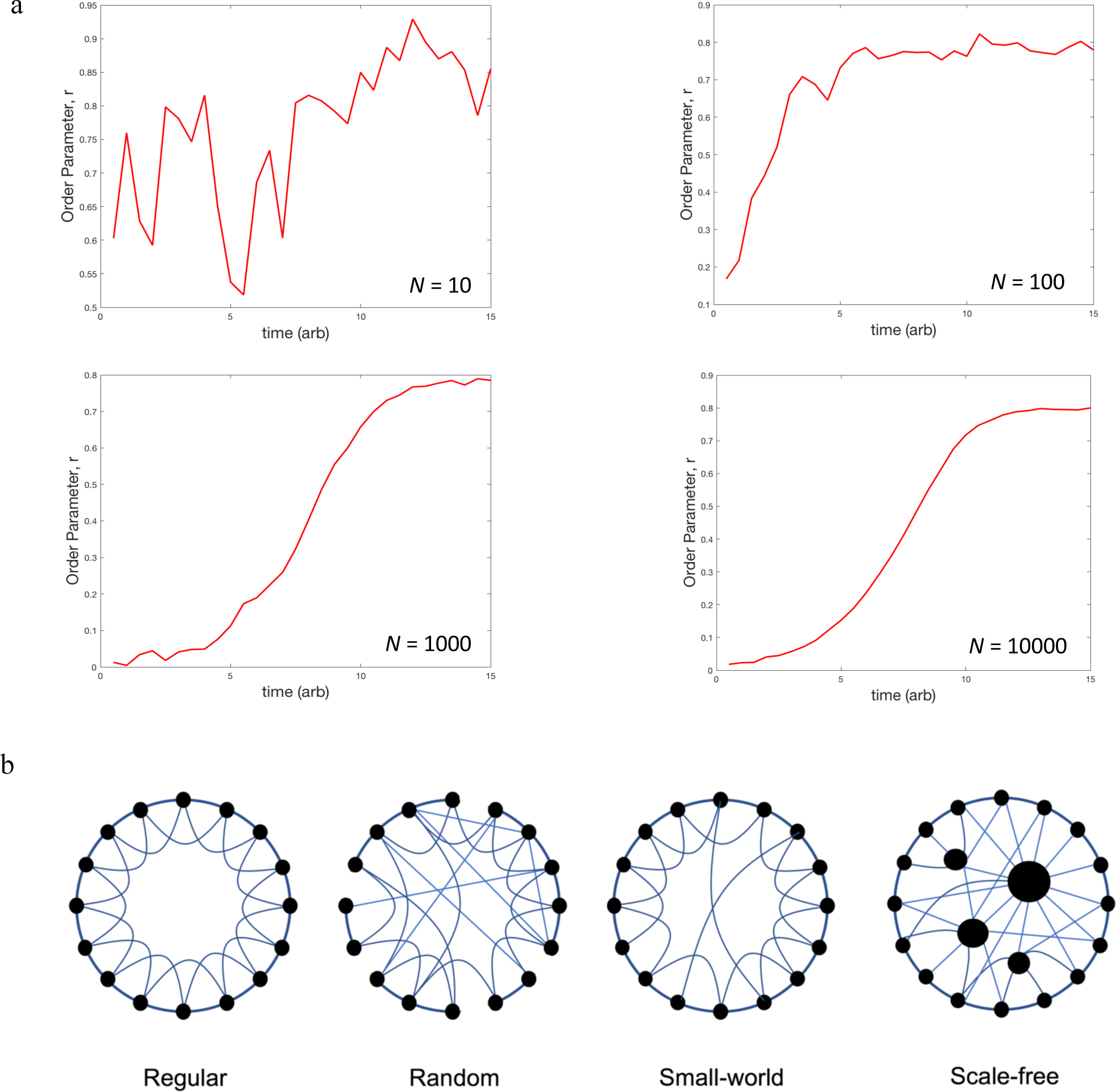
Synchronization and Networks. a) Increasing coupled oscillators leads to synchronization for a fixed coupling strength. Order parameter, *r*, against time show transition from unordered to smooth ordered response over time (arbitrary units) as coupled oscillators, *N*, is increased for a fixed coupling strength, *K* = 1.9. Note that numerous simulations for each *N* were performed since each run will give different kinetics, nevertheless, the general response for each *N* is indicative of plot shown. b) Regular, Small-world, Random and Scale-Free Networks.

As mentioned earlier, one area of biology that faces the challenge of *sync* behavior is the development of resistance mechanisms of microorganisms, such as bacteria, through biofilm formation that evades drug treatment [6]. Here, the study to *desync* the emergent collective behaviors could benefit the development of effective therapeutics.

## 3. Small World and Scale Free Networks favor *sync*

In a 1998 paper, Watts and Strogatz investigated the collective dynamics in regular, random and small-world networks (Fig. 1B) [15]. They also devised an order parameter (*p* = 0 being order and *p* = 1 is disorder) based on characteristic path length and clustering coefficient ratios. Very importantly, without investigating complex dynamics, their methodology applied on static (steady-state) data showed that information propagation and order behavior is spread more effectively in small-world networks rather than the other forms of networks.

Subsequent works investigated the dynamics of small-world networks using Kuramoto oscillators [16-18]. It was shown that phase *sync* emerged even for a small number of short cuts, which was otherwise not possible for regular or random networks with the same conditions [16]. A similar result was also obtained when using a different (Rossler) type of oscillators on small-world networks [19]. In other words, *sync* is easier to achieve in small-world network rather than random networks. Thus, do biological systems, which has shown *sync* behavior on numerous situations, tap the benefit of “short cuts” or small-world networks?

It is now well-known that biological networks are large and organized in a scale-free manner [20]. Although scale-free network also displays short cuts, the main difference between small-world and scale-free networks is that the latter possess a relatively low number of hubs; nodes that are highly connected to other nodes (Fig. 1B). The distribution of the nodes and their connectivity follows a power-law, hence the name scale-free [20]. Moreno and Pacheco (2004) tested this network using Kuramoto oscillators and found similar result to the small-world networks, that *sync* is more optimally achieved even for low coupling strength [21]. Thus, the size and architecture of coupled oscillators in a network play a key role in *sync*, and that the scale-free network observed in biology seems to be a key feature for collective self-organized response.

## 4. Biological Response - large invariance in low and middle expressed genes

As mentioned earlier, biological networks, such as transcriptional networks, are known to possess scale-free property [12]. We recently investigated transcriptome-wide expressions data of *Saccharomyces cerevisiae* biofilm, in wildtype and 6 different single-gene overexpressed strains (*SAN1, TOS8, ROF1, SFL1, HEK2, DIG1*), related to biofilm regulation using various statistical methods [22]. These strains were previously studied due to their biofilm morphological regulation [23].

Our analysis revealed that for any single gene overexpression strain, despite producing noticeable local expression changes by the 296 acute genes (Fig. 2A, orange dots), the global transcriptome-wide structure (constructed by the majority of 2,486 collective genes) was not perturbed in any significant manner (Fig. 2A, black dots). This indicates that there is strong global regulatory transcriptome-wide structure or invariance. Hence, over time, the biofilm morphological changes generated by each overexpression strain will most likely be attenuated.

**Figure 2.**
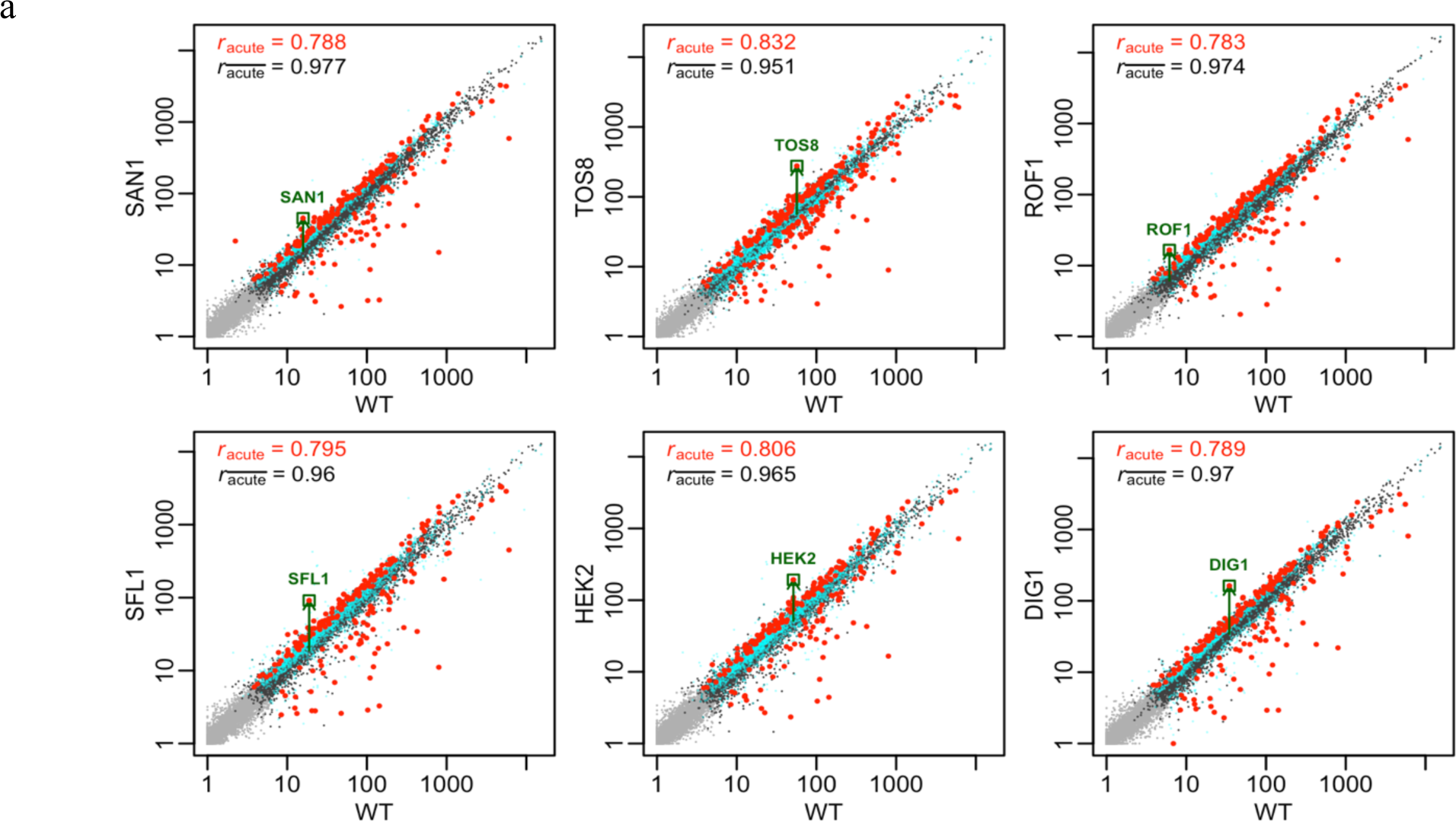
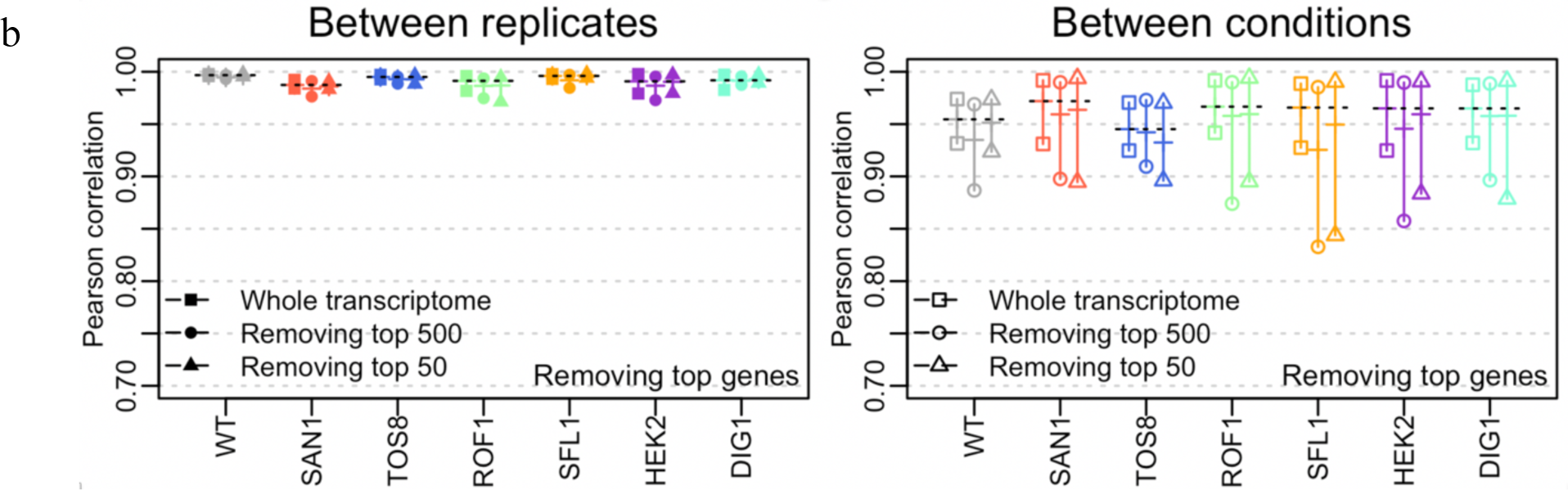
Strong transcriptome-wide correlation between different yeast biofilm strains. a) comparing gene expressions between wildtype and different biofilm regulation strains shows invariance for the large proportion of genes and b) Pearson correlation between replicates (left) and strains (right) for each strain is high (mean > 0.9) despite removing 50 or 500 highest expressed genes. Figure reused from [22].

Notably, the high transcriptome-wide correlation between the different strains were mainly attributed to the low to middle expressed genes. That is, out of over 11,000 transcript expressions, we found the top 50-100 expressed genes were poorly correlated between the different strains. This result was confirmed even by comparing the correlation with random extraction of 50-100 genes from whole transcriptome [22]. Furthermore, removing the top 50 or 500 expressed genes from the data still showed correlation close to unity (Fig. 2B). Gene ontology analysis suggested that many of the low to middle expressed genes constituted important biological functions such as nucleotide metabolism, cell cycle or stress response. This result is consistent with Figure 1A, where the Kuramoto model predicts that for a large network, like the transcriptional network, low coupling strength (or expressions) are sufficient to generate synchronization.

Based on the study, we, therefore, postulate that the low and middle expressed genes, comprising of diverse cellular processes, are necessary for global cell population to be in *sync*. Thus, biofilm control using single gene target may not be a viable option for successful long-term effects as the large majority of genes are unchanging and tightly controlled (strong order parameter)

Future work could track their measurements more closely in time to test the application of Kuramato model for biological *sync*. However, the challenge is that these genes may not be reliably tested at individual expression levels due to the interference of technical noise on their relatively low expression levels [24-25].

## 5 Conclusion

In this paper, I have highlighted the concepts of *sync* using the example of the popular Kuramoto model. Examples on the type of network that favors *sync* are also mentioned: both small-world and scale-free networks are able to *sync* at lower coupling strength compared to regular or random networks. To compare the significance in biology, which possess scale-free networks, our transcriptome-wide analysis of several strains of yeast biofilm show that low to middle expressed genes are very highly correlated (even when small sampling of genes was performed) between the different genotypes, an indication of biological *sync*. Further work requires measuring the correlated genes’ expressions reliably over time and find target(s) that would *desync* the transcriptome-wide collective response in cases where biofilm is a nuisance, such as for antibiotic resistance.

## 6 Acknowledgments

The author thanks BioTrans, A*STAR, for supporting the publication of this article.

